# A seed resource for screening functionally redundant genes and isolation of new mutants impaired in CO_2_ and ABA responses

**DOI:** 10.1101/415976

**Authors:** Felix Hausera, Paulo H. O. Ceciliatoa, Yi-Chen Lin, DanDan Guo, JD Gregerson, Nazia Abbasi, Diana Youhanna, Jiyoung Park, Guillaume Dubeaux, Eilon Shani, Nusra Poomchongkho, Julian I. Schroeder

## Abstract

The identification of homologous genes with functional overlap in forward genetic screens is severely limited. Here we report the generation of over 14,000 amiRNA-expressing plants that enable screens of the functionally redundant gene space in *Arabidopsis*. A protocol is developed here for isolating robust and reproducible amiRNA-mutants. Examples of validation approaches and essential controls are presented for two new amiRNA mutants that exhibit genetically redundant phenotypes and circumvent double mutant lethality. In a forward genetic screen for abscisic acid (ABA)-mediated inhibition of seed germination, amiRNAs that target combinations of known redundant ABA receptor and *SnRK2* kinase genes were rapidly isolated, providing a strong proof of principle for this approach. A new ABA insensitive amiRNA line is isolated, which targets three genes encoding *avirulence-induced gene2-like* (*AIG2*) genes. A thermal imaging screen for plants with impaired stomatal opening in response to low CO_2_ exposure led here to isolation of a new amiRNA targeting two essential proteasomal subunits, PAB1 and PAB2. The seed library of 14,000 T2 amiRNA lines generated here provides a new platform for forward genetic screens and is being made available to the Arabidopsis Biological Resource Center (ABRC) and optimized procedures for amiRNA screening and controls are described.

**Highlight:** The generation of over 14,000 amiRNA-expressing plants is reported that are being made publicly available enabling screens of redundant genes in *Arabidopsis*. Identification of known and new genes is reported.

## Introduction

The presence of large gene families in plants including *Arabidopsis thaliana* (Arabidopsis Genome Initiative, 2000) leads to functional genetic redundancies or partial functional overlap among closely related genes. Functional overlap and partial or complete redundancy between different family members has been proposed to provide a buffer for loss or gain of function mutation events and mechanistic robustness of cellular networks (Wagner, 2005). This is considered to be a main reason for the lack of observable phenotypes in single-gene deletion mutants and increasing severity of phenotypes in higher order mutants of homologous genes (Ma *et al.*, 2009; Park *et al.*, 2009). Identification and characterization of functionally overlapping genes in genetic screens is limited, as is evident by the relatively low number (591 of all *Arabidopsis* genes) of genes not associated with a single mutant phenotype (Lloyd and Meinke, 2012). Analysis of genome-wide gene family definitions showed that the *Arabidopsis* genome includes over 22,000 genes belonging to gene families (Hauser *et al.*, 2013). Strategies and tools have been developed to enable screens of the functionally redundant gene space. Recently, an artificial microRNA (amiRNA) based computational design approach was introduced (Hauser *et al.*, 2013). AmiRNAs designed to specifically target diverse combinations of gene family members or combinations of subfamily members enable the screening of partial over-lapping homologous gene functions at a genome-wide scale. The presented platform also provides an approach for capture of homologous gene silencing phenotypes, for which higher order loss of function mutants would lead to lethality, as illustrated by a mutant identified here.

Here we report the generation of over 14,000 T2 amiRNA lines by transformation of *A. thaliana* Col-0 with a previously published amiRNA library (Hauser *et al.*, 2013) and screening of T2 amiRNA lines for abscisic acid (ABA) insensitive seed germination phenotypes or plants with low CO_2_ insensitive high leaf temperature phenotypes. Methods are described to identify robust amiRNA mutants and how to avoid pitfalls of this approach. The screen rapidly identified two amiRNAs which target three *PYR/RCAR ABA* receptors (Ma *et al.*, 2009; Park *et al.*, 2009) or six *SNF1-related kinase* (SnRK2s (Mustilli, 2002; Yoshida *et al.*, 2002; Fujii and Zhu, 2009)) encoding genes known to be involved in ABA-mediated control of seed germination. One candidate line which shows an ABA insensitive seed germination phenotype contains an amiRNA that targets three genes of unknown function which are annotated as AIG2A (AT3G28930), AIG2LA (AT5G39720) and AIG2LB (AT5G39730). One amiRNA which causes a low CO_2_ insensitive high leaf temperature phenotype targets two genes encoding proteasomal !2-subunits annotated as PAB1 (AT1G16470) and PAB2 (AT1G79210), for which double mutation causes lethality. New amiRNA lines that target the proteasomal **α**([0-9]+)-subunit annotated as PAG1 (AT2G27027), was constructed resulting in a similar stomatal phenotype. Together these observations indicate a rate-limiting role of the intact proteasome for stomatal opening responses.

## Materials and Methods

### Plant material, growth conditions and transformation

*Arabidopsis thaliana* accession Columbia-0 was used as the background for all amiRNA transformations of the library. Surface sterilized seeds (15 min 70% Ethanol, 0.1% Sodium dodecyl sulfate; 3-4 washes with ∼ 100% Ethanol; alternatively 10 min 50 % bleach, 0.05% Tween 20; 4 - 6 washes with water (Lindsey *et al.*, 2017) of *A. thaliana* were cold-treated for 2 - 5 days at 4°C and germinated on half strength Murashige and Skoog basal medium supplemented with Gamborg’s vitamins (Sigma-Aldrich (Murashige and Skoog, 1962; Gamborg *et al.*, 1968), 0.8% Phytoagar (Difco, Franklin Lakes, NJ) and pH adjusted (pH 5.8; 4-Morpholinoethane sulfonic acid (2.6 mM; Sigma-Aldrich) titrated with potassium hydroxide). After 5 – 7 days, plants were transferred to plastic pots containing sterilized premixed soil (Sunshine Professional Blend LC1 RS; Sunshine; supplemented with an appropriate amount of insecticide (Marathon, Gnatrol)) and propagated under the following conditions: long day (16hr light/ 8 hrs. dark); 23-27 °C; 20-70% humidity, 60-100 mmol m^−2^ sec^−1^ light.

Plant transformation by floral dip was performed as described elsewhere (Clough and Bent, 1998) with the following modifications: *Agrobacterium tumefaciens* GV3101::pMP90 (Koncz and Schell, 1986) was grown under selection of all markers, i.e. genomic (rifampicin), Ti-plasmid (gentamicin), pSOUP (tetracycline) and T-DNA plasmid (spectinomycin). The infiltration medium for resuspension of the bacteria and floral dip contained 5% Sucrose (w/v) and 0.02% (v/v) Silwet L-77 (Clough and Bent, 1998).

Large scale transformation with the amiRNA library pools (Hauser *et al.*, 2013) was performed as described elsewhere (Cutler *et al.*, 2000) with the following modifications. One microgram DNA of each amiRNA sub library (Hauser *et al.*, 2013) was electroporated into a total of 500 μl electrocompetent *A. tumefaciens* cells. The 20 bp and 21 bp amiRNA sub library variants for each pool were individually electroporated and combined at this stage. After two hours at 30 °C in non-selective Luria Bertani Miller medium (LB, Teknova) the cells were spread on 20 LB plates (1.5% agar; 150 mm diameter) containing all the appropriate antibiotics (rifampicin, gentamycin, tetracycline, spectinomycin) and grown for three days at 30°C. The bacteria were scraped from the plates, re-suspended in 5 ml LB and concentrated by centrifugation for 20 min at 5855 *x g*. Plants were transformed by spraying the flowers with this suspension of the bacteria in infiltration medium (adjusted to an optical density at 600 nm of 0.5) twice with one week between the treatments. T1 plants were selected on plates supplemented with 75 μM phosphinotricin or directly on soil by spraying diluted herbicide (1000 x dilution, Finale^®^; Bayer, North Carolina Research Triangle Park, NC) four times with 2 - 7 days between the treatment. Herbicide resistant plants were transferred to soil and grown to full maturity and T2 seeds collected from individual plants. When appropriate, media for growth of bacteria or plant selection contained the following concentrations of antibiotics (mg ml^−1^): carbenicillin 100, gentamycin 25, kanamycin 30, rifampicin 50, spectinomycin 100, tetracycline 10, and phosphinotricin 15.

### Screen for abscisic acid insensitive seed germination phenotype

T2 plants were screened individually for insensitivity of seed germination to abscisic acid in 96 well plates (100 μl 0.1 % Agarose supplemented with 2 μM (±)-ABA, Sigma-Aldrich). Approximately 10 - 20 seeds were used from each T2 plant. For the pooled screening, approximately 10 - 50 seeds of 90 individual T2 plants were mixed, surface sterilized and sprinkled onto agar plates (3 μM (±)-ABA; Sigma-Aldrich). As control for ABA insensitivity, *abi4-1* (ABRC, CS8104) or *abi5-1* (ABRC, CS8105) were used, Col-0 was used as wild-type control. A putative abscisic acid (ABA) insensitive phenotype was scored in a binary manner for similarity to the *abi4-1* or *abi5-1* phenotype and difference to wild type after 5 - 8 days using green cotyledons as indicator (Kuhn *et al.*, 2006). For lines which showed a putative ABA insensitivity, the seed germination assay was repeated by propagating individual T2 seedlings to the next generation (T3) and using seeds of the T3 generation for ABA sensitivity assays. This time, seeds were placed on plates with and without ABA (2 μM (±)-ABA; Sigma-Aldrich) and images were taken daily for 7 days and emergence of radicles and cotyledons was counted manually using Fiji (Schindelin *et al.*, 2012). For candidates of the individual screen the T2 seeds were used for the repetition of the germination assay.

For candidates of the pooled screen ABA insensitive seedlings were transferred to plates containing 75 μM phosphinotricin and after 7 – 10 days resistant seedlings were transferred to soil, grown up to full maturity and the T3 seeds used for the validation of the ABA insensitive germination phenotype.

### Screen for CO_2_ insensitive leaf temperature phenotype

Seeds of T2 plants were germinated in 96 pots flats (254 mm x 508 mm; East Jordan Plastics, East Jordan, MI) on soil with each pot containing seeds from one plant. After seven days seedlings were sprayed with a 1000 x dilution of Finale^®^ (Bayer, Bayer, North Carolina Research Triangle Park, NC) and two to three days later pale seedlings were removed and only one healthy dark green seedling was left per pot. After 19 days under standard growth conditions the plants were exposed to 150 ppm CO_2_ for two hours in a Percival growth chamber. A first set of thermal images was taken with a FLIR A320 thermal imaging camera (FLIR, Wilsonville, OR). Subsequently the plants were exposed to ≥ 800 ppm CO_2_ and after two hours a second set of thermal images was taken. Control plants included in the experiments were *ht1-2* (Hashimoto *et al.*, 2006), *ost1-3* and wild-type Col-0. Thermal images were converted into Flexible Image Transport System format (fits) using the ExaminIR software (FLIR, Wilsonville, OR). For the screen using the 96-pot flat format, the temperature of plant leaves and the surrounding soil were measured using Fiji (Schindelin *et al.*, 2012). The soil temperature served as location specific reference to compensate for temperature variation depending on the position in the 96 pots flat. Either the temperature difference between plant leaves and surrounding soil or the average temperature of plant leaves were used as a quantitative measure. Plants with more than one degree Celsius difference to soil were considered as primary candidates and subject to further testing. The high-temperature leaf phenotype of *ht1-2* was used as a reference for CO_2_ insensitivity. To test the reproducibility of the CO_2_-dependent leaf temperature phenotype of putative candidates, T2 plants were grown in triplicate and assayed again alongside with *ht1-2* and wild-type control plants.

### Identification of amiRNA sequences and testing reproducibility

Genomic DNA from candidates with a robust and reproducible phenotype was isolated as described elsewhere (Edwards *et al.*, 1991) and the sequence of the amiRNA present was determined by sequencing of the PCR product (primers pha2804f and pha3479r; see Supplemental Table 1). Using the Target Search function available on the WMD3 website (Ossowski *et al.*, 2008) putative amiRNA target genes were predicted. For independent confirmation of the phenotype independent lines were generated by cloning the identified amiRNA into pFH0032 (Supplemental Table 2; (Hauser *et al.*, 2013) and transforming it into *Arabidopsis* Col-0. Confirmed phenotypes were further analyzed by using single knock out mutants, higher order mutants generated by crossing and/or generating amiRNAs which target subsets of the initial target genes (Supplemental Table 3).

### Gas exchange analyses

Stomatal conductance of H_2_O (gs) was measured in leaves of 5 to 6-week-old plants using portable gas exchange systems (LI-6400 and LI-6800, LI-COR, Lincoln, Nebraska), starting 2 hours after growth chamber light onset. For intact single leaf ABA treatments, a LED light source set at 150 µmol m^−2^ s^−1^ (10% blue) and a chamber temperature of 21 °C was used. Leaves were equilibrated for one hour at a relative humidity of 70-72%, airflow of 200 rpm and CO_2_ concentration of 400 ppm. After one-hour, steady-state stomatal conductance was recorded ten minutes before the addition of ABA to the petioles in water at the indicated concentration. For light-response measurements, plants were kept in the dark for 18 hours prior to experiments. Stomatal conductance of a single intact leaf in the dark was recorded for 10 min, followed by red light treatment of 600 μmol m^−2^ s^−1^. After 20 minutes of red light treatment, additional blue light was applied at 10 μmol m^−2^ s^−1^. The incoming air humidity was kept at 62-65% and air flow at 200 rpm. For stomatal conductance measurements of single intact leaf CO_2_-responses, incoming relative air humidity was kept at 62-65% and the imposed changes in CO_2_ concentration were applied as indicated. Leaves were attached to intact plants and were equilibrated for one hour before the measurements. The data presented represent n ≥ three leaves with each leaf from independent plants per genotype per treatment.

### qRT-PCR analysis

Total RNA (500ng) was reverse transcribed using the first-strand cDNA synthesis kit (GE Healthcare). qRT-PCR analyses were performed using 3-fold-diluted cDNA, Maxima SYBR Green Rox/qPCR Master Mix (Thermo Scientific). The housekeeping *PDF2* gene was used as an internal control. The threshold cycle (CT) was determined by the instrument, and the Δ ΔCT method was used to calculate the fold change in each gene (Livak and Schmittgen, 2001). For *RAB18* gene expression measurements, two-week-old seedlings were treated with ABA for nine hours, final concentration of 20 μM and total RNA extraction was used.

## Results and Discussion

### Generation of amiRNA library plants

We have previously described the generation of an amiRNA library consisting of ten sub libraries that represent 22,000 individual amiRNA designs (Hauser *et al.*, 2013). Deep sequencing of these 10 sub libraries showed that Δ 95% of the designed amiRNAs were present in these sub libraries (Hauser *et al.*, 2013). The amiRNA library was transformed first into *Agrobacterium tumefaciens* and then into *Arabidopsis* Col-0. Over a period of over four years, the amiRNA library consisting of ten sub libraries was transformed and T1 seeds harvested. Using plate or soil-based selection methods, herbicide resistant T1 plants were grown and T2 seeds from over 14,000 individual plants were harvested (Table 1). The transformation rate varied over a range from 0.08% to 0.76% with an average of 0.25%. During the course of this research, approximately 3,000 additional T2 lines were generated expressing amiRNAs that target homologous transporter-encoding gene family members. These 3,000 lines will also be made available to the ABRC, such that over 14,000 total T2 lines will be submitted for use by the community.

**Table 1.** Overview of the ten amiRNA libraries as described by Hauser et. al. (2013), the number of amiRNAs designed for each library and the number of individual T2 amiRNA transformants that were generated here. Note that for the generation of each pool the 20bp and 21bp libraries were combined (see Hauser et. al. 2013 for details). All T2 lines have been submitted to the Arabidopsis Biological Resource Center (ABRC).

| **Pool name** | **Pool description** | **amiRNA designs** | **T2 lines** |
| --- | --- | --- | --- |
| kinase receptor (PKR) | Protein kinases, protein phosphatases, receptors and their ligands | 1860 | 817 |
| binding (BNO) | Proteins binding small molecules | 1968 | 1771 |
| protein (CSI) | Proteins that form or interact with protein complexes including stabilization of those | 2313 | 842 |
| rna dna (TFB) | Transcriptionfactors and other RNA and DNA binding proteins | 2964 | 831 |
| metabolism (TEC) | Metabolic and other enzymes catalyzing transfer reactions (EC: 2) | 1521 | 1548 |
| diverse enzymes (PEC) | Catalytic active proteins, mainly enzymes | 1881 | 1113 |
| non-classified (UNC) | Genes for which the function is not known or cannot be inferred | 4082 | 1387 |
| diverse molfun (DMF) | Protein with diverse functional annotation not found in the other categories | 1505 | 1152 |
| hydrolase (HEC) | Hydrolytic enzymes (EC class 3), excluding protein phosphatases | 2129 | 971 |
| transporter (TRP) | Proteins that transport molecules across membranes | 1777 | 3844 |
| TOTAL |  |  | 14276 |

### Screen for ABA insensitive seed germination phenotype

In total over 2,500 T2 amiRNA lines were screened individually and over 5,000 T2 amiRNA lines were screened in pools for ABA insensitive germination phenotypes (Figure 1). In the primary screen using individual plants in a 96-well plate format, 59 putative candidates were identified. In the primary screen using pools of 90 plants with 25 - 80 seeds per line, 340 putative candidates representing an unknown number of lines were identified (Figure 1).

**Figure 1.**
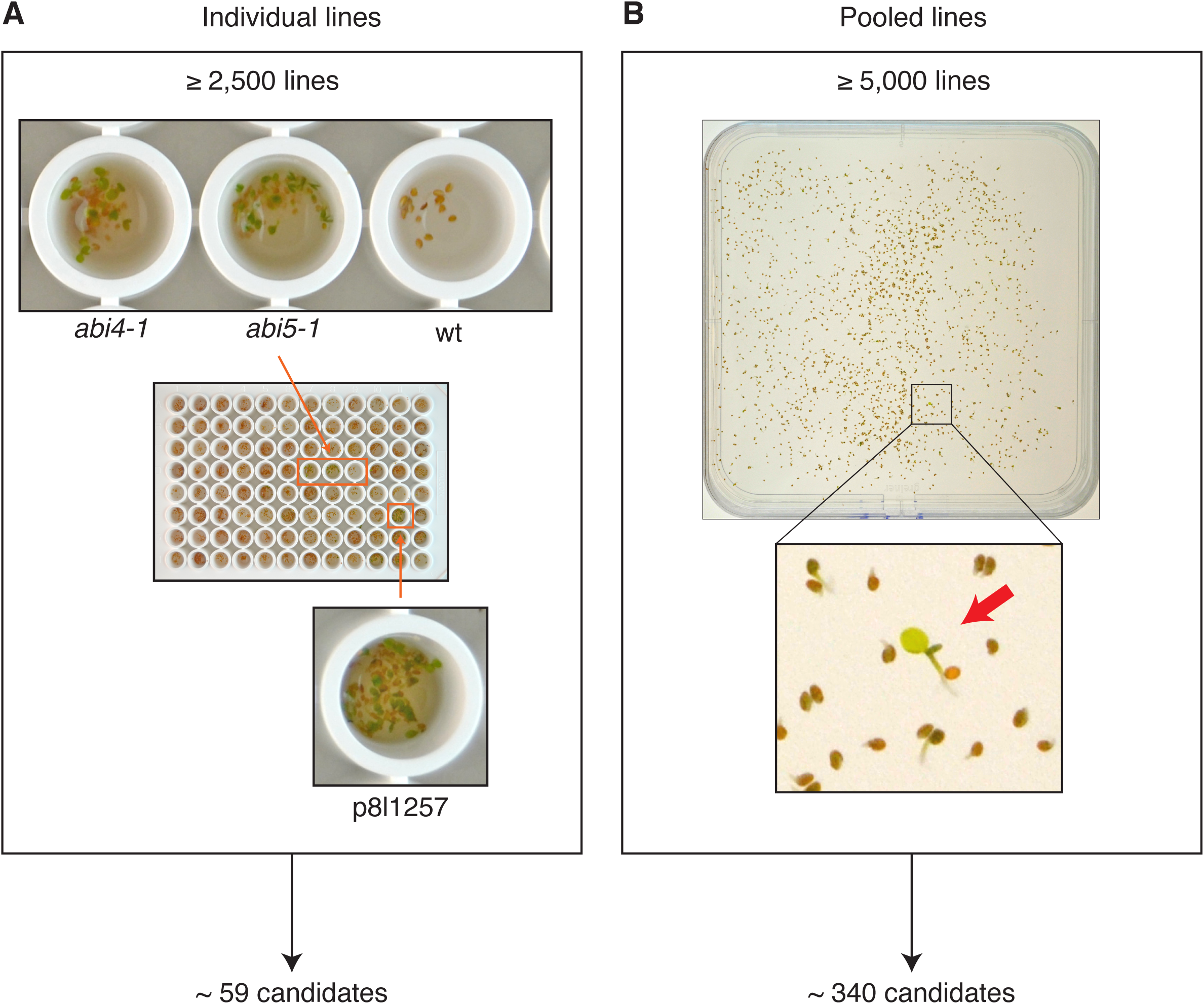
Overview of primary screen performed with over 7,500 T2 amiRNA lines. **(A)** Approximately 2,500 T2 amiRNA lines were screened individually by adding seeds to 96 well plates each well containing 100 μl 0.1% Agarose supplemented with 2 μM ABA and approximately 10 – 20 seeds from one plant. Wild type (Col-0), *abi4-1* and *abi5-1* were used as controls. Based on visual inspection of cotyledon greening around 59 lines were considered as candidates for further testing in the T3 generation. **(B)** Approximately 5,000 T2 amiRNA lines were screened in pools. Each pool contained 10 – 50 seeds from 90 individually stored amiRNA lines (see text). Approximately 340 lines were selected for further testing in the T3 generation.

These candidates were subjected to further analysis in a secondary screen (Figure 2). The cotyledon emergence phenotype of 24 T3 seedlings from a total of 76 retested plants showed a more reduced ABA sensitivity that was clearly different from wild type and less severe than the abi4-1 and abi5-1 controls (Figure 2A). From the 59 putative candidates identified using the individual screening approach, the amiRNA line p8l1257 showed a reproducible partial insensitivity to ABA in the T3 generation (Figure 3). Only the amiRNA in candidates with the most robust phenotypes were determined by sequencing. Two of the amiRNA-targeted gene sets identified in 24 seedlings with reproducible phenotypes are known core components of the ABA signal transduction cascade (Figure 2, Table 2). These include amiRNA lines that target the three ABA receptors PYR1 (RCAR11), PYL4 (RCAR10) and PYL6 (RCAR9) (Figure 2B, C and Table 2). Furthermore, amiRNA-expressing plants that target six members of the SnRK2 protein kinase family (Mustilli, 2002; Yoshida *et al.*, 2002; Fujii and Zhu, 2009) were isolated in this screen, including the three SnRK2 protein kinases, SnRK2.2, SnRK2.3 and SnRK2.6 (OST1) that are known to be required for abscisic acid signaling (Figure 2B, C and Table 2; (Mustilli, 2002; Yoshida *et al.*, 2002; Fujii and Zhu, 2009)).

**Figure 2.**
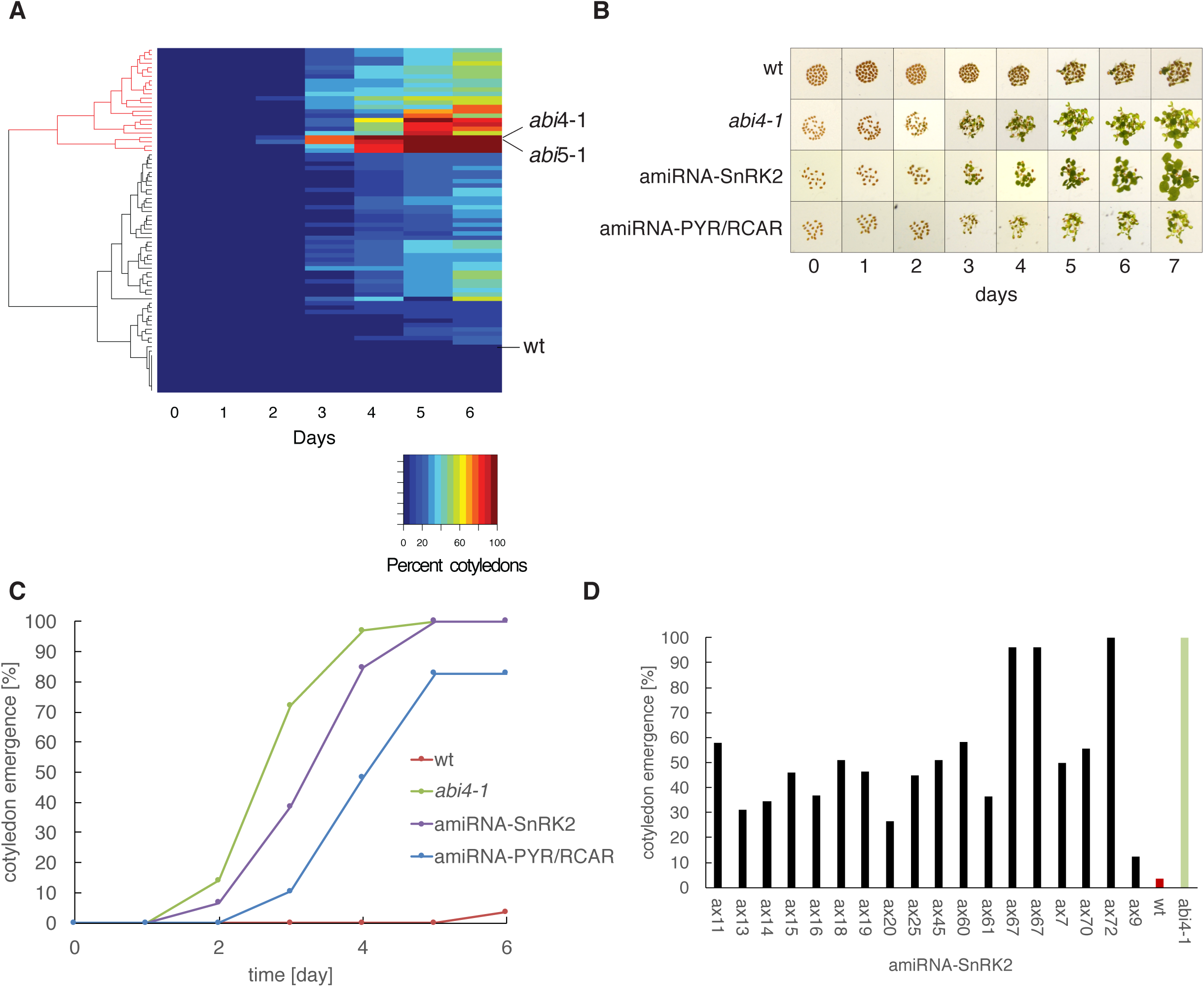
Overview of secondary ABA insensitivity screen performed with T3 candidate amiRNA lines identified in the primary screen which target known genes involved in ABA signal transduction. **(A)** Heat map representation of cotyledon emergence time course of the T3 generation obtained from candidates with putative ABA insensitive seed germination or cotyledon emergence phenotypes. Each row represents percent cotyledon emergence of one individual line. The rows are ordered by hierarchical clustering. Wild-type control (WT, Col-0) and *abi4-1* and *abi5*-1 as reference for ABA insensitivity are shown. **(B)** Images of seeds germinating on plates in secondary screen containing 2 μM ABA at the indicated time points. Shown are representative plants of amiRNA plants targeting a set of six *SnRK2* kinase genes (*amiRNA-SnRK2*) or a set of three *PYR/RCAR* ABA receptor genes (*amiRNA-PYR/RCAR*; see Table 2). Wild-type control (Col-0) and *abi4-1* and *abi5-1* as reference for insensitivity are shown in the other two rows. **(C)** Time course of hypocotyl emergence in the presence of ABA for wild-type control (wt), *abi4-1* as a reference for ABA insensitivity and two representative amiRNA lines that target a set of six *SnRK2* protein kinase genes (*amiRNA-SnRK2*) or a set of three *PYR/RCAR* ABA receptor genes (*amiRNA-PYR/RCAR*). Approximately 74 ± 46 seeds were phenotyped per genotype. **(D)** Variation of cotyledon emergence phenotype (day six; 2 μM ABA) in the T3 generation of plants isolated in the primary screen which were selected as candidates based on their seed germination phenotype in the T2 generation. Sequencing of the amiRNA confirmed that all 18 of these amiRNA expressing plants contain an amiRNA that targets a set of six *SnRK2* kinase genes (*amiRNA-SnRK2*; see Table 2). Wild-type control (wt, Col-0) and *abi4-1* as reference for ABA insensitivity are shown.

**Figure 3.**
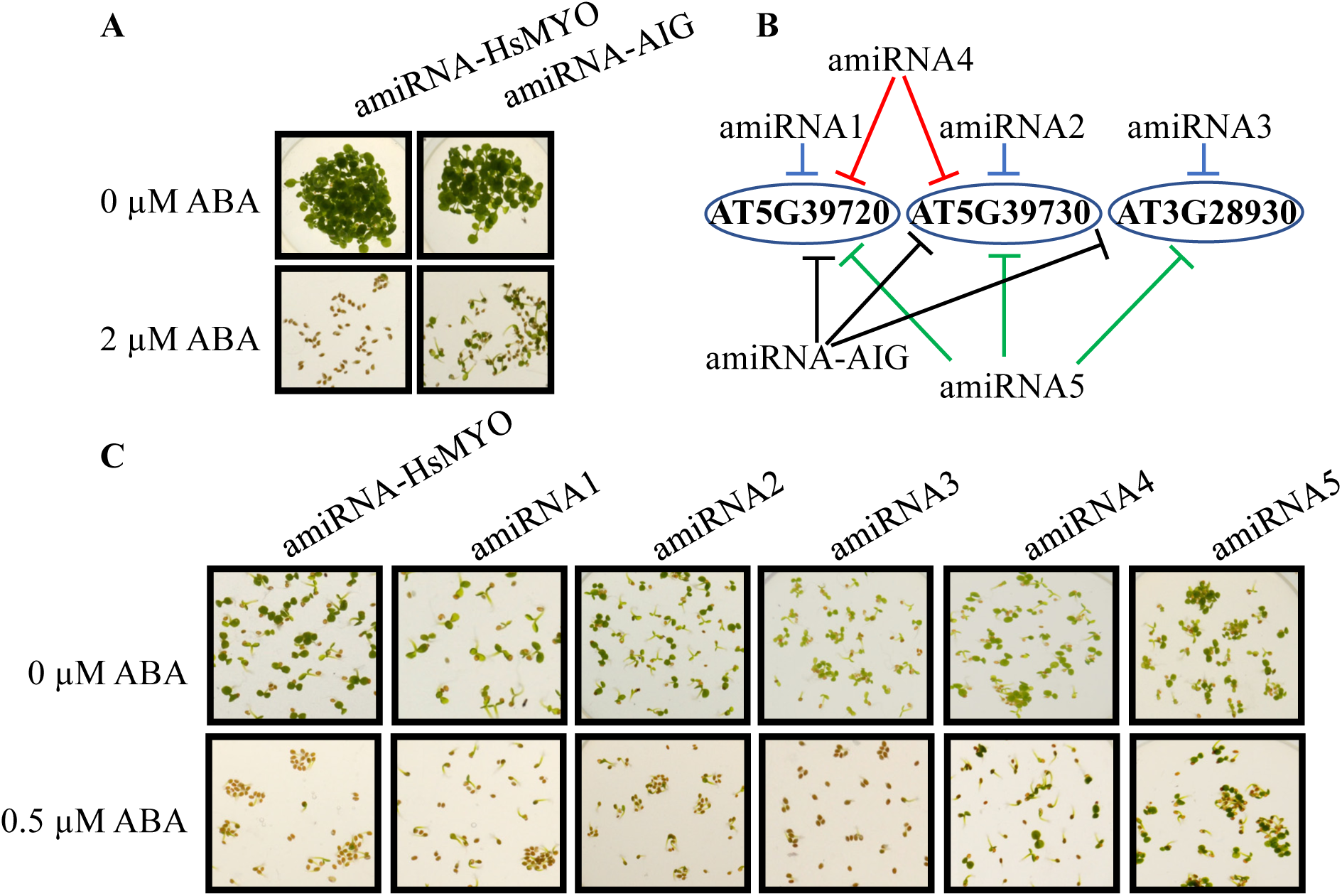
Avirulence-Induced Genes (AIGs) targeted by an amiRNA cause a reduced ABA sensitivity in cotyledon emergence assays. **(A)** Seedlings of the control amiRNA line, which has no target gene in *Arabidopsis* (*amiRNA-HsMYO* control) and an amiRNA line which targets three *AIG* genes (*amiRNA-AIG*) germinated in the presence of 0 or 2 μM ABA. Photographs were taken after 12 days of exposure. **(B)** Five new amiRNAs were designed to target the three AIG genes. Three of these amiRNAs, amiRNA1, 2 and 3, target a single gene each. AmiRNA4 targets two tandem-repeat *AIG* genes and amiRNA5 targets all three genes at non-identical nucleotides compared to the original amiRNA isolated in the primary screen (*amiRNA-AIG*, see Supplemental Table 2 for amiRNA sequences). **(C)** The new T2 amiRNA lines were tested in cotyledon emergence assays. Seedlings were germinated in the presence of 0 or 0.5 μM ABA. Photographs were taken after 4 days of incubation.

**Table 2.**
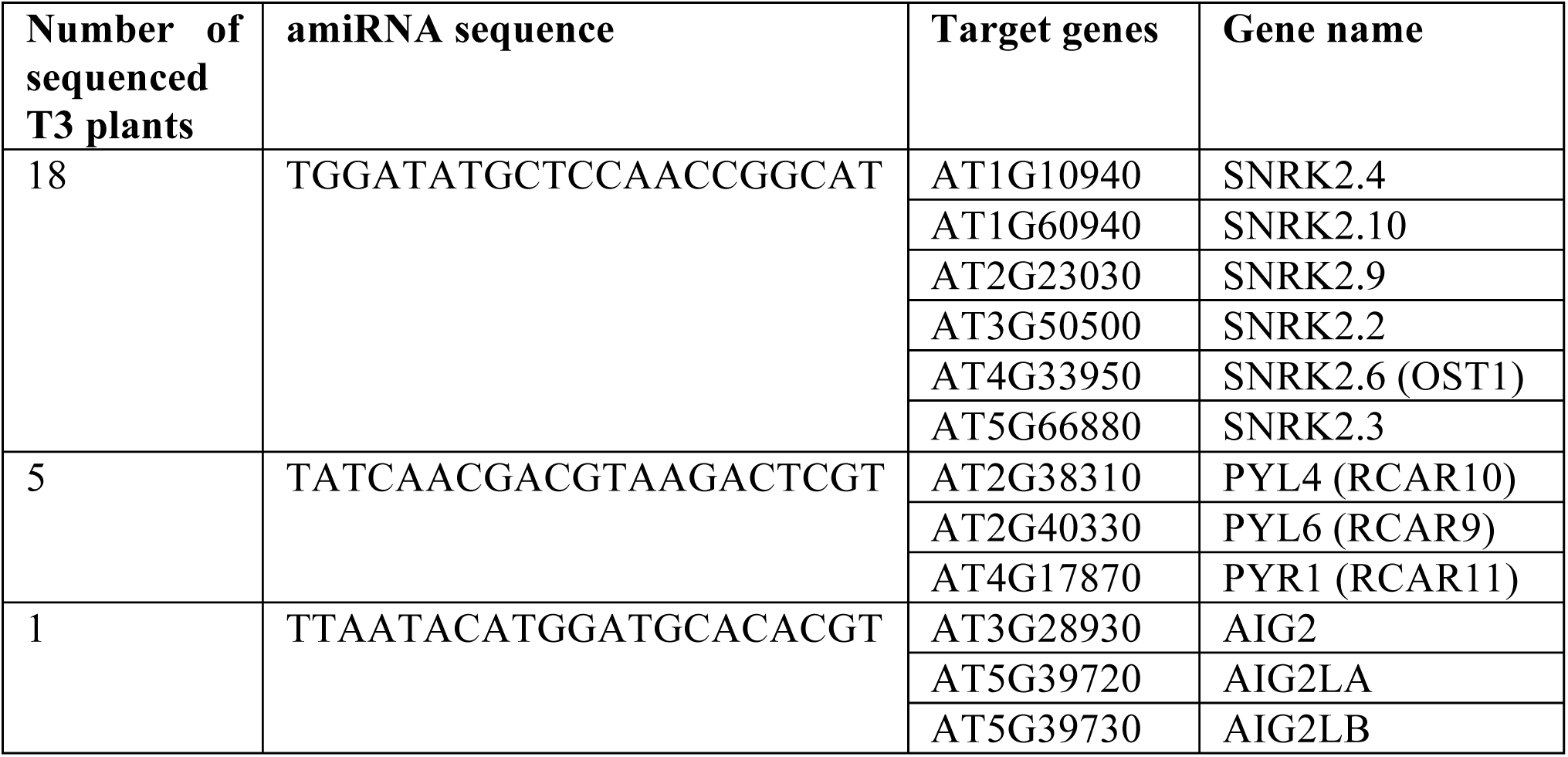
AmiRNA sequences and predicted target genes found in candidate T3 plants showing robust ABA insensitive seed germination phenotypes in the T2 screen and subsequently in the T3 generation.

Notably, Figure 2D shows a strong variation in the cotyledon emergence phenotype among plants expressing the same amiRNA that targets six *SnRK2* kinase transcripts. This variation might be responsible for the high number of variable candidates which did not show a robust phenotype following the primary screen. Additional amiRNA lines were isolated as putative mutants and the amiRNA sequence was determined (Supplemental Table 4). Although some of the predicted targets might be expected to affect abscisic acid responses, rescreening of these putative mutants did not show consistently robust reproducible phenotypes. Thus, amiRNAs appear to produce phenotypes that may be variable even within the same line. These findings led us to develop a protocol in which:
1. Only putative mutants that showed a consistent phenotype when screening seeds from the next generation of plants were selected.
2. Only lines that showed similar phenotypes upon re-transformation with new amiRNAs that are predicted to target the same genes were selected. Furthermore, based on the variation observed here in the secondary screen it is advisable to investigate over ten independent transformed lines (Schwab *et al.*, 2006; Hauser *et al.*, 2013) in the future to determine which amiRNAs produce phenotypes that can be carried forward. The isolation of amiRNA lines targeting functionally over-lapping *PYR/RCAR* ABA receptor and *SnRK2* protein kinase genes, that would not be isolated in traditional forward genetic mutant screens, provides a proof of principle that functionally redundant genes can be isolated in forward genetic screening using this new amiRNA resource. The inclusion of control lines and the validation steps described above should enable screening for diverse phenotypes using the lines generated here that are being provided to ABRC.

### AmiRNA lines targeting three avirulence-induced genes (*AIGs*) show partial insensitivity to ABA-inhibition of seed germination but not to ABA-induced stomatal closure

The amiRNA in line p8l1257 isolated in the present screen targets a new set of three genes (Figure 3A and 3B). Previous research annotated these genes based on their mRNA upregulation in a transcriptomic study after infection with *Pseudomonas syringae pv maculicola* carrying avrRpt2 (<u>a</u>vrRpt2-induced <u>g</u>ene, *AIG2*) (Reuber and Ausubel, 1996). However, these genes have not been previously described to be involved in ABA-mediated control of seed germination or other phenotypic responses in plants.

The line p8l1257 was named *amiRNA-AIG* here and was further tested by analyzing seed germination with additional T2 generation seeds from the original p8l1257 stock (Figure 3). Germination properties were compared to a control amiRNA line targeting the human myosin 2 (*amiRNA-HsMYO*), that has no targeted genes in *Arabidopsis* (Hauser *et al.*, 2013). After 12 days on plates containing 2 μM ABA, the *amiRNA-AIG* line showed cotyledon greening in contrast to the control *amiRNA-HsMYO* line (Figure 3A). The effect of the amiRNA-AIG on the expression of a known ABA induced gene *RAB18* was analyzed by qRT-PCR (Figure S1). The ABA mediated induction of *RAB18* expression was substantially reduced in the *amiRNA-AIG* line indicating a role of the targeted AIG genes in ABA signal transduction.

Since two out of the three genes are tandemly repeated, generation of double mutants using T-DNA insertion knockouts would be limited. Therefore, five new amiRNA lines were generated which target subsets of genes targeted by the original *amiRNA-AIG* to verify the relevance of the predicted *AIG* target genes. *AmiRNAs 1, 2* and *3* targeted each a single *AIG* (Figure 3B; Supplemental Table 3). *AmiRNA 4* targeted two tandem-repeat *AIG* genes and *amiRNA 5* targeted all three *AIG* genes targeted in the original *amiRNA-AIG* line, but with a different amiRNA sequence (Figure 3B, see Supplemental Table 3 for amiRNA sequences). When the T2 seeds expressing these five new amiRNAs were tested in a seed germination assay with 0.5 μM ABA, only the *amiRNA 4* and *amiRNA 5*-expressing lines showed less sensitivity to ABA compared to the control *amiRNA-HsMYO* line in cotyledon greening (Figure 3C). The expression of all three putative target genes (AT5G39720, AT5G39730, AT3G28930) was analyzed using qRT-PCR in the originally isolated *amiRNA-AIG* line and in all the *amiRNA* lines 1 to 5 (Figure S2). The amiRNA efficiency of transcriptional inhibition varies between the lines, target transcript(s) and amiRNA sequence. Note that microRNA silencing in plants occurs via two mechanisms, (a) the degradation of transcripts and (b) inhibition of translation (Brodersen *et al.*, 2008). Thus, quantification of targeted transcripts may not fully show the degree of silencing of target genes. Combined, these data provide evidence that the original *amiRNA-AIG* phenotype is attributable to silencing of more than one *AIG* gene, suggesting overlapping homologous gene functions.

The original *amiRNA-AIG* line was also investigated to determine if it affects ABA-induced stomatal closure using an intact leaf gas exchange analysis approach. When ABA was applied to the transpiration stream of intact leaves at a final concentration of 2 μM, both the control *amiRNA-HsMYO* line and the *amiRNA-AIG* lines showed an ABA-induced decrease in stomatal conductance to H_2_O (gs, Figure 4A). Normalization of the stomatal conductance data showed no dramatic difference in ABA-induced stomatal closure between *amiRNA-HsMYO* and *amiRNA-AIG* (Figure 4B). Together, the present data show that the isolated *amiRNA-AIG* line is less sensitive to ABA-inhibition of seed germination.

**Figure 4.**
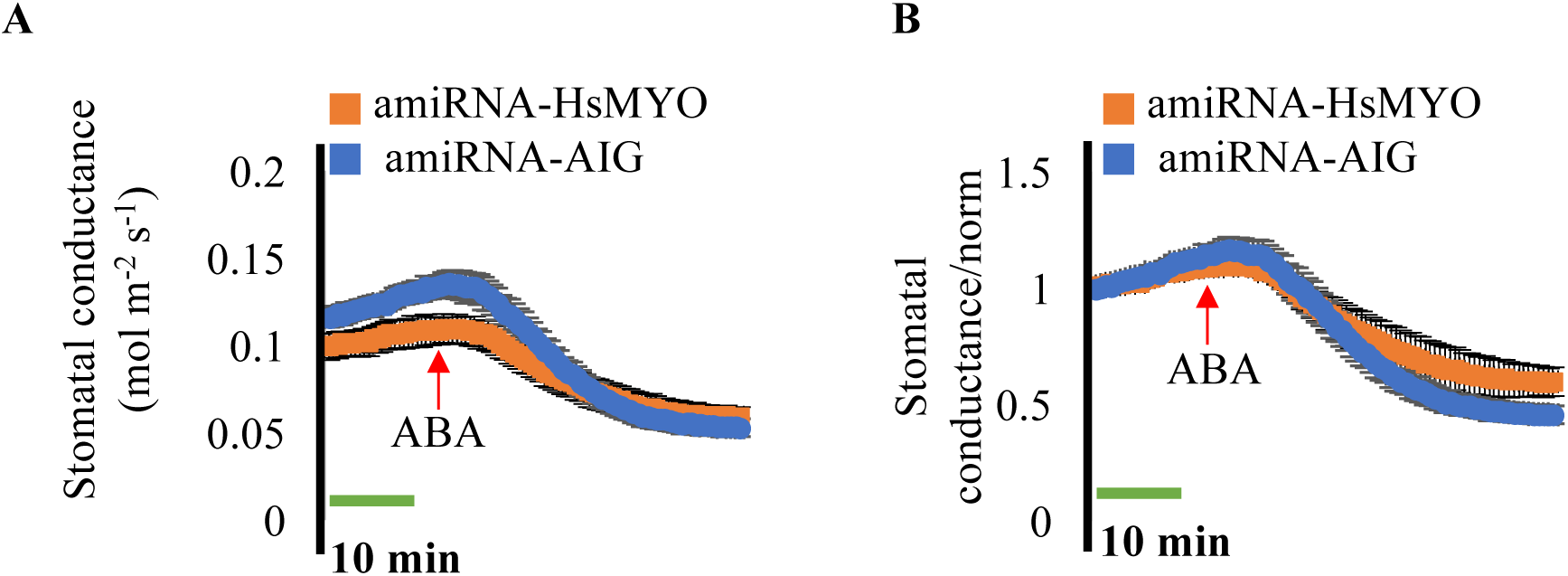
The isolated amiRNA line targeting three *AIG* genes (*amiRNA-AIG*) responds to ABA in whole leaf gas exchange analyses. Time-resolved stomatal conductance to H_2_O (gs) in response to application of 2μM ABA to the transpiration stream (red arrows) are shown in the amiRNA control line, (*amiRNA-HsMYO*) and in the amiRNA line targeting three *AIG* genes (*amiRNA-AIG*). Stomatal conductance was analyzed using whole leaf gas exchange. **(A)** Stomatal conductance in mol m^−2^s^−1^. **(B)** Data from **(A)** were normalized to stomatal conductance at the beginning of the experiment. Data are the mean of n = 3 leaves per genotype ± S.E.M.

The *AIG2* genes are functionally annotated as putative γ-glutamyl cyclotransferases (GGCTs, EC:4.3.2.9) based on their similarity to the human orthologue (HsGGCT; O75223). AIG2LA and AIG2LB share only 16% and 17% identity respectively to the human orthologue. GGCTs have been described to cleave γ-glutamyl-amino acid dipeptides to release the free amino acid and 5-oxoproline (Oakley *et al.*, 2008). Further research will be required to determine the mechanism by which AIG2s affect ABA inhibition of seed germination.

### Screen for CO_2_ insensitive leaf temperature phenotype

In total, over 2,500 T2 amiRNA lines were screened individually for an altered leaf temperature response to a low CO_2_ concentration (150 ppm) by infra-red thermal imaging (Figure 5). Leaf temperature depends on various parameters including radiation absorption, air temperature and humidity (Merlot *et al.*, 2002). Low ambient CO_2_ concentration leads to stomatal opening in *Arabidopsis*, causing an increased transpiration rate and thus a decrease in leaf temperature compared to the surrounding air. Mutants impaired in CO_2_-induced stomatal opening appear warmer compared to wild type plants. In the screen, we used the soil temperature as reference to compensate for the local temperature differences due to various factors including humidity of the soil. Wild-type plants and plants of the HIGH TEMPERATURE1-2 (*ht1-2*) mutant (Hashimoto *et al.*, 2006) were included in all trays as controls. Based on visual inspection of the thermal images, plants with relatively higher leaf temperature under low [CO_2_] compared to the other plants in the same image were selected and the difference between the average leaf temperature and the surrounding soil was determined. The difference between leaf temperature and soil temperature was determined as reference for overall temperature and to compensate for local temperature differences. A set of 106 plants with more than one degree difference between the leaf temperature and the surrounding soil was defined as initial putative candidates for further testing (see Methods for details). For the rescreening of putative mutants, we set a high threshold for temperature differences in the selection of mutants compared to the wild-type strain of 1°C. The constitutive CO_2_ response mutant *ht1-2*, when exposed to low [CO_2_], shows a delta temperature above 1 °C between leaf and soil. Rescreening of these candidates in the T2 generation revealed an amiRNA line (*p9l22*) with a robust and reproducible impaired response to low CO_2_ (Figure 5B).

**Figure 5.**
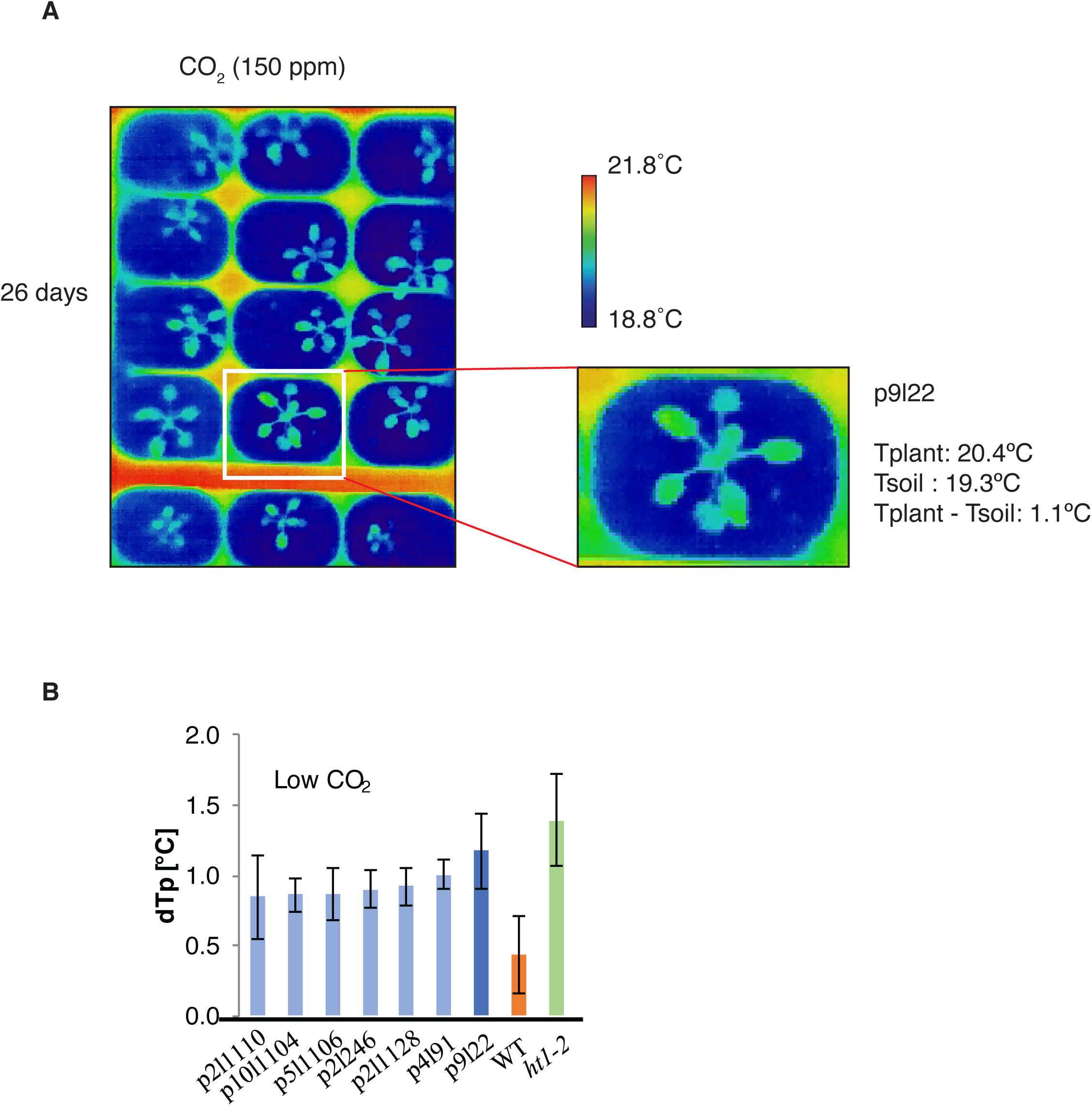
Thermal imaging screen for mutants with impaired response to low CO_2_ identified the amiRNA line p9l22 **(A)** Thermal images of amiRNA lines in the primary screen after exposure to low CO_2_ (150ppm). Plants were grown in 96 pot flats under ambient CO_2_ and after 26 days exposed to low (150 ppm) CO_2_ for two hours and then thermal images of the entire flat were taken in 8 separate images per flat. An image is shown from the primary screen in which the plant *p9l22* (white box) was flagged as a candidate with a putative altered response to CO_2_ based on the leaf temperature. The average leaf temperature was computationally calculated across all rosette leaves for flagging putative mutants. **(B)** Differences of leaf temperature (Tplant) and surrounding soil (Tsoil) for putative mutants exposed to low [CO^2^] (150 ppm) for two hours. Average leaf temperature was computed by image analysis of the most fully expanded rosette leaves. Bars show average ± SD (n=3 to 5 leaves). WT (Col-0) (orange) and *ht1-2* (green) were used as control.

After exposure to low [CO_2_], the leaf temperature of the *p9l22* line was compared to wild type (Col-0) and to the constitutive high CO_2_-response mutant *ht1-2* (Figure 6A; (Hashimoto *et al.*, 2006)). The leaves of the *p9l22* line had a higher temperature than wild-type leaves and a similar temperature to *ht1-2* leaves (Figure 6A). Stomatal index (SI) and density (SD) were calculated for wild type, the control *amiRNA-HsMYO* and *p9l22* lines. No noteworthy differences were found between the genotypes (Figure S3; *amiRNA-HsMYO* vs. *p9l22* line, One-Way ANOVA, p-value >0.05 for SI and SD).

**Figure 6.**
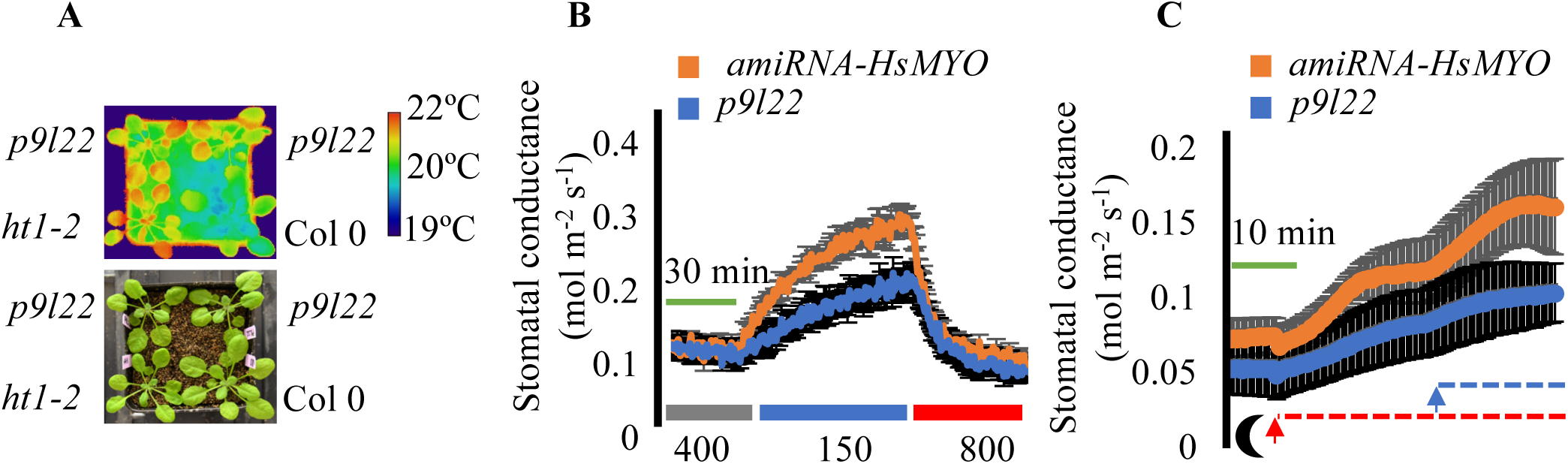
The amiRNA line *p9l22* is defective in light and low CO_2_-induced stomatal opening. **(A)** The *p9l22* line shows an elevated leaf temperature phenotype at low ambient [CO_2_] treatment (150 ppm). Thermal imaging: (top) and image of the same plants: (bottom). The calibration bar shows the pseudo-colored scale for temperature. **(B)** Time-resolved stomatal conductance responses at the imposed [CO_2_] shifts (bottom in ppm) in the control *amiRNA-HsMYO* line and in the *p9l22* line were analyzed using intact whole-leaf gas exchange. The plots represent average ± S.E.M (n= 4 leaves from 4 plants per genotype). **(C)** Time-resolved stomatal conductance responses from darkness to the imposed light intensity and light quality shifts (bottom moon shape: darkness; red dashed line: red light at 600 µmol m^−2^ s^−1^ and blue dashed line: blue light at 10 µmol m^−2^ s^−1^) in the control *amiRNA-HsMYO* line and in the *p9l22* line were analyzed using intact whole-leaf gas exchange. The plots represent average ± SE (n= 3 leaves from 3 plants per genotype).

To measure [CO_2_] responses in a time-resolved fashion, we measured stomatal conductance (*g*_*s*_) using a gas exchange analyzer. In the *amiRNA-HsMYO* control line, the shift from ambient (400 ppm) to low (150 ppm) [CO_2_] led to a rapid increase in stomatal conductance (Figure 6B). *AmiRNA* line *p9l22* responded to the same treatment with a lower magnitude of stomatal opening (Figure 6B). Both lines showed stomatal closure in response to high (800 ppm) [CO_2_] exposure at similar rates (Figure 6B). To test whether line *p9l22* is defective in response to other stimuli that cause stomatal opening, light-induced *g*_*s*_ responses were investigated (Figure 6C). The control *amiRNA-HsMYO* and *p9l22* lines were kept in the dark for 18 hours prior to the experiments and steady-state *g*_*s*_ was measured. When red light (at 600 μmol m^−2^ s^−1^) was applied, the *p9l22* line showed a reduced rate of *g*_*s*_ increase when compared to the control line. The same was observed when blue light (at 10 μmol m^−2^ s^−1^) was superimposed on the red light background (Figure 6C). Thus, the amiRNA line *p9l22* causes reduced responses to low CO_2_ concentration, red light and blue light.

The amiRNA in the *p9l22* line was sequenced and is predicted to target two closely homologous proteasomal subunit genes (*PAB1,* At1G16470 and *PAB2*, At1G79210). Both *PAB1* and *PAB2* genes are the sole two genes that encode the 20S proteasome alpha 2 (**α**([0-9]+)) subunit (Baumeister *et al.*, 1998). First, we attempted to isolate a double mutant (*pab1 pab2*) using T-DNA insertion lines (SALK_099950 and SALK_144987) (Alonso *et al.*, 2003). After genotyping over 100 plants in the F2 generation, no homozygous double mutant was recovered. We concluded that the double mutation is very likely to be lethal.

Alternatively, a new amiRNA sequence targeting solely the *PAB1* gene was cloned and transformed into the *pab2-1* single mutant (SALK_144987). This new amiRNA line, *pab2-1*mut *pab1amiRNA*, was investigated in stomatal conductance analyses of [CO_2_] responses (Figure 7). Leaves were first exposed to high (900 ppm) [CO_2_] for one hour and steady-state *gs* was recorded. Shifts to low (150 ppm) [CO_2_] led to an increase in *gs* in both the *pab2-1*mut *pab1amiRNA* line and the control *HsMYO-amiRNA* line (Figure 7A). The normalized stomatal conductance data show that *pab2*mut *pab1amiRNA* line responds to low [CO_2_] with a reduced magnitude compared to the control line (Figure 7B).

**Figure 7.**
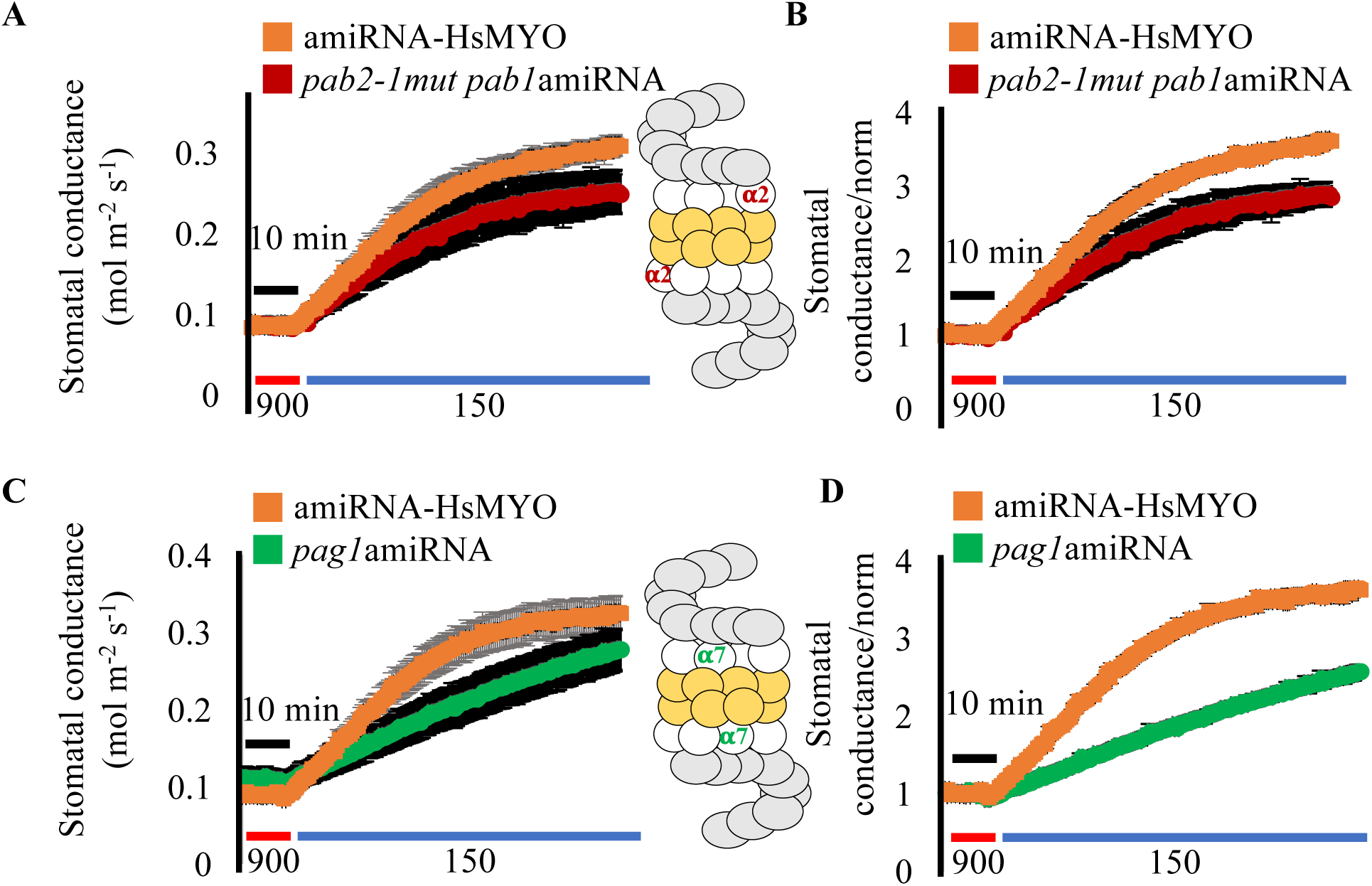
New amiRNA lines targeting the two *PAB* genes (**α**([0-9]+) subunit) and *PAG1* gene (**α**([0-9]+) bunit) of the 26S proteasome show partial impairment in low CO_2_-induced stomatal opening. Time-resolved stomatal conductance responses at the imposed [CO_2_] shifts (bottom in ppm) in the control line *amiRNA-HsMYO* and **(A)** in the *pab2-1mut pab1amiRNA* line (*pab2* gene T-DNA knockout and *pab1* gene amiRNA knockdown, **α**([0-9]+) subunit) and **(C)** *pag1amiRNA* (*pag1* gene amiRNA knockdown, **α**([0-9]+) subunit) line were analyzed using intact whole-leaf gas exchange. The plots represent average ± SE (n= 3 to 4 leaves from different plants each/genotype). **(B and D)** Data from **(A)** and **(C)** were normalized to the average stomatal conductance of the first 10 minutes. Inserts show a representation of the 26S proteasome, with the **α**([0-9]+) subunits highlighted in red and **α**([0-9]+) subunits highlighted in green. The initial slope of gs response for *amiRNA-HsMYO* and *pag1amiRNA* was calculated and One-Way ANOVA test was used to compare the values (p-value > 0.05).

Initial experiments were pursued to determine if modifications in the alpha ring of the 20S proteasome might be linked to the above phenotypes, or whether this mutation is specific to only !2 subunit mutations of the proteasome. The alpha ring of the 20S proteasome is composed of seven alpha subunits (Kurepa and Smalle, 2008). The *p9l22 amiRNA* targets the only two genes that encode the !2 subunit of the proteasome (Figure 7, inset highlighted in red). To determine whether other alpha subunits also affect the response to low [CO_2_], a second amiRNA line was generated, which targets the *PAG1* gene (**α**([0-9]+) subunit, inset in Figure 7 highlighted in green), named *pag1*amiRNA. The **α**([0-9]+) subunit is encoded by a single gene in *Arabidopsis* (Kurepa and Smalle, 2008). When a *pag1*amiRNA line was tested in *gs* responses to [CO_2_] shifts, it showed a lower rate of stomatal opening when compared to the control *amiRNA-HsMYO* line (Figure 7C, D; One-Way ANOVA p-value >0.05). The expression levels of *PAB1, PAB2* and *PAG1* were analyzed in the *p9l22* amiRNA line and also in *pab2-1*mut *pab1amiRNA* and *pag1amiRNA* lines using qRT-PCR (Figure S4). With the exception of the severely reduced *PAB2* expression in the *pab2-1*mut *pab1amiRNA* line when compared to control lines, no clear evidence for knock down at the transcriptional level could be detected in the amiRNA lines, which may point to amiRNA mediated inhibition of translation (Brodersen *et al.*, 2008).

The present findings show that the *p9l22* amiRNA line is also partially impaired in red light-induced stomatal opening. Red light mediates stomatal opening in part via activation of photosynthesis and the resulting drop in internal concentration of CO_2_ (Ci) (Roelfsema *et al.*, 2002; Matrosova *et al.*, 2015). In addition, the *p9l22* line is also partially impaired in blue light-induced stomatal opening. This suggests that a general regulator of stomatal opening is impaired in this line. As the proteasome mediates the degradation of proteins and reduced functions of α-ring subunits are predicted to increase protein levels, it is tempting to speculate that the phenotype observed might be correlated with an increased abundance of a negative regulator of stomatal opening. Further research will be required to test this or other hypotheses. In other studies, the 26S **α**([0-9]+) subunit, when overexpressed, enhanced thermotolerance and adaptation in rice and *Arabidopsis*, suggesting that proteasomal subunits can have rate-limiting roles in regulating plant physiological responses (Li *et al.*, 2015).

### Summary and Future Use

A library of over 14,000 total T2 generation amiRNA lines was created as a new resource to screen the redundant gene space in *Arabidopsis*. These amiRNA-expressing lines are being provided as individual lines to the Arabidopsis Biological Resource Center (ABRC). Given the observations and findings in the present study, lines will be available for high-throughput screening in pools of 90 lines per pool with approximately 25 to 50 seeds per individual amiRNA line in each pool. In each pool, the pooled seeds for screening will originate from one of the 10 sub-libraries that target gene family members with defined functional annotations (Table 1; (Hauser *et al.*, 2013)). This approach will increase the probability of identifying interesting putative mutants in future screens despite the biological variability in amiRNA silencing lines found here (Figure 2D).

The screen for abscisic acid insensitive seed germination phenotypes identified two amiRNAs targeting *PYR/RCAR* ABA receptor genes and *SnRK2* genes, that are both known groups of redundant key genes and proteins required for ABA signal transduction (Ma *et al.*, 2009; Park *et al.*, 2009). Isolating amiRNA lines in these known components serves as proof of principle for our approach. Moreover, screening this amiRNA population enables the identification of mutants that require co-silencing of homologous gene family members, which are less likely to be found in forward genetic screens of EMS or T-DNA mutagenized seed populations. Overall the presented amiRNA screen shows that amiRNA lines are prone to show a high rate of variable candidates with weak or non-robust phenotypes. Nevertheless, as shown here new mutants can be isolated. Furthermore, during the course of this research, this amiRNA library resource has been used to isolate long-sought functionally redundant auxin transporter genes (e.g. *ABCB6, ABCB20*; Zhang *et al.,* 2018). Approaches to circumvent the inherent limitations of forward genetic screening with amiRNAs are developed in the present study. As a first step, it is recommended to rescreen the next generation to identify robust and reproducible phenotypes in individually isolated putative mutant lines. As a second step, the amiRNA sequence of confirmed mutant lines needs to be determined (see Methods). AmiRNA sequences linked to the observed phenotypes are retransformed and testing over 10 independent lines for the phenotype is recommended. Alternatively, amiRNAs which target a subset of the initially predicted targets can be used to narrow down the causative genes (e.g. Figure 3). For cases where only two to three genes are targeted T-DNA lines or CRISPR/Cas9 mutants can be used to narrow down the genes relevant for the phenotype.

Over 95% of the amiRNAs in this library were designed to target only two to five genes (Hauser *et al.*, 2013), meaning that identification of causal genes is facilitated. Using the above approach, we report on two newly identified mutants: (i) amiRNA lines targeting three genes encoding avirulence induced gene 2 like proteins show an ABA insensitive seed germination phenotype. (ii) amiRNA lines targeting two proteasomal subunits show insensitivity to low CO_2_ induced stomatal opening. Further analyses of the two targeted genes in this amiRNA line, suggest that stronger T-DNA alleles result in lethality. This indicates the usefulness of the generated amiRNA lines for forward genetic isolation of higher order mutants that would be lethal upon knock out. Our data suggest that the wild-type expression levels of two alpha subunits of the 20S proteasome, **α**([0-9]+) and **α**([0-9]+), are required for fully functional stomatal opening mediated by physiological stimulation. This indicates that the proteasomal subunits are likely controlling an unknown general negative regulator of stomatal opening. The amiRNA seed resource generated here provides a new and potent tool to identify redundant genes and also lethality causing higher order mutants in many biological processes in Arabidopsis.

In conclusion, the amiRNA library resource is well-suited for screening of phenotypes that can be easily verified in subsequent generations. This population may be best suited for screens that permit high throughput or medium throughput screening for phenotypes with a large dynamic range. Many such screens have been performed in classical *Arabidopsis* mutant screens that were not designed to identify functionally redundant genes.

## Supplementary data

**Supplemental Table 1**. List of relevant primers used in this study.

**Supplemental Table 2**. List of relevant plasmids used in this study.

**Supplemental Table 3**. List of new amiRNAs designed and cloned in this study.

**Supplemental Table 4.** AmiRNA sequences and predicted target genes found in candidate plants.

**Figure S1.** The induction of *RAB18* gene expression by ABA is lower in the amiRNA-AIGs line.

**Figure S2.** The expression of *AIGL* genes in amiRNA-AIGs lines.

**Figure S3.** The *p9l22* amiRNA line has normal stomatal indices and density.

**Figure S4.** The expression of *PAB1, PAB2* and *PAG1* genes in amiRNA lines.

## Acknowledgements

Seeds for the over 14,000 T2 amiRNA lines described here are being made available to the Arabidopsis Biological Resource Center (ABRC) for screening by the research community (order numbers: CS99427, CS99428, CS99429, CS99430, CS99431, CS99432, CS99433, CS99434,CS99435, CS99436). We thank Dr. Jianyan Huang, Kellie Tao Kim, Wilma Lee, Sandra Vogel, Marianne Kreusch and Elly Poretsky for help in the transformation of the amiRNA library into *Arabidopsis*, and generation of the lines and support during the various stages of the screen for novel phenotypes. This research was funded by grants from the National Science Foundation (MCB1616236) and the National Institutes of Health (GM060396-ES010337) to J.I.S. and was in part supported by grants from the Israel Science Foundation (1832/14) and a European Research Council Starting Grant (757683 - RobustHormoneTrans) to E.S. P.H.O.C was supported by a Ciencias sem Fronteiras/CNPq fellowship. G.D. was supported by an EMBO long-term postdoctoral fellowship (ALTF334-2018).

